# Expression of *Ace2, Tmprss2*, and *Furin* in mouse ear tissue

**DOI:** 10.1101/2020.06.23.164335

**Authors:** Tsukasa Uranaka, Akinori Kashio, Rumi Ueha, Taku Sato, Han Bing, Gao Ying, Makoto Kinoshita, Kenji Kondo, Tatsuya Yamasoba

## Abstract

**Objectives:** Intracellular entry of the severe acute respiratory syndrome coronavirus 2 (SARS-CoV-2) depends on the interaction between its spike protein to a cellular receptor named angiotensin-converting enzyme 2 (ACE2) and depends on Furin-mediated spike 23 protein cleavage and spike protein priming by host cell proteases including 24 transmembrane protease serine 2 (TMPRSS2). *Tmprss1, Tmprss3,* and *Tmprss5* are expressed in the spiral ganglion neurons and the organ of Corti in the inner ear; however, *Ace2, Tmprss2,* and *Furin* expression profiles in the middle ear remain unclear. Therefore, this study aimed to analyze *Ace2, Tmprss2,* and *Furin* expression in the middle and inner ear of mice.

**Study Design:** Animal research.

**Setting:** Department of Otolaryngology and Head and Neck Surgery, University of Tokyo.

**Methods:** We performed immunohistochemical analysis to examine the distribution of Ace2, Tmprss2, and Furin in the eustachian tube, middle ear space, and cochlea of mice.

**Results:** Ace2 was expressed in the cytoplasm in the middle ear epithelium, eustachian tube epithelium, stria vascularis, and spiral ganglion. Tmprss2 and Furin were widely expressed in the middle ear spaces and the cochlea.

**Conclusion:** Co-expression of Ace2, Tmprss2, and Furin in the middle ear indicates that the middle ear is susceptible to SARS-CoV-2 infections, thus warranting the use of personal protective equipment during mastoidectomy for coronavirus disease (COVID-19) patients.

## Introduction

The recent coronavirus disease (COVID-19) pandemic caused by severe acute respiratory syndrome coronavirus 2 (SARS-CoV-2) poses a serious health concerns. SARS-CoV-2 generally infects the upper respiratory tract and then spreads to various organs. The clinical symptoms of COVID-19 patients include fever, cough, sore throat, fatigue, and loss or decline of smell (anosmia or hyposmia).^1–4^ The Eustachian tube is the conduit between the middle ear space and upper respiratory tract through which, SARS-CoV-2 can spread into the middle ear spaces. Moreover, other human coronaviruses have been identified in middle ear fluid in cases of acute otitis media or otitis media with effusion.^5–7^ Two studies have reported the occurrence of acute otitis media^8^ and sensorineural hearing loss^9^ among COVID-19 patients. Another study carried out pure tone audiometry and transitory evoked otoacoustic emission (TEOAE) analysis for 20 asymptomatic patients aged 20–50 years positive for COVID-19 on reverse transcription-PCR (RT-PCR) testing and reported that high-frequency pure-tone thresholds and TEOAE amplitudes were significantly worse among the COVID-19 patients than among 20 healthy controls.^10^

Intracellular entry of SARS-CoV-2 depends on the interaction between viral spike proteins and a cellular receptor, angiotensin-converting enzyme 2 (Ace2)^11–13^, Furin-mediated Spike cleavage^13–15^, and Spike protein priming by host cell proteases including transmembrane protease serine 2 (Tmprss2).^13,16^ Thus, Ace2, Tmprss2, and Furin upregulation enhances intracellular SARS-CoV-2 entry, thus resulting in clinical symptoms. Previous studies have reported 8 *Tmprss* genes including *Tmprss2* in the inner ear through RT-PCR analysis, although immunohistochemical analysis revealed the expression of only Tmprss 1/3/5 in the inner ear, indicating that Tmprss 1/3/5 are expressed in the spiral ganglion neurons and Tmprss3 is also expressed in the organ of Corti.^17^ To our knowledge, Ace2, Tmprss2, and Furin expression in the middle ear spaces and Ace2 and Furin expression in the inner ear have not been reported.

This study aimed to investigate the mechanism underlying middle ear infection and sensorineural hearing loss in COVID-19, by assessing *Ace2*, *Tmprss2*, and *Furin* expression in the middle and inner ear tissues of mice.

## Materials and Methods

### Experimental samples

Tissue samples were obtained from the mice used in our previous studies because the purchase of new animals has been prohibited in our facility owing to the COVID-19 pandemic. Tissue samples were obtained from normal 11-week-old male ICR mice, and paraffin-embedded tissue samples including the middle ear spaces, Eustachian tube area, and cochlea were used (Figure 1, 2). Normal tissue morphology in these regions was confirmed through hematoxylin and eosin staining by a qualified pathologist and by otolaryngologists. All experiments were conducted in accordance with institutional guidelines and with the approval of the Animal Care and Use Committee of the University of Tokyo (No. P18-015)

**Figure 1.**
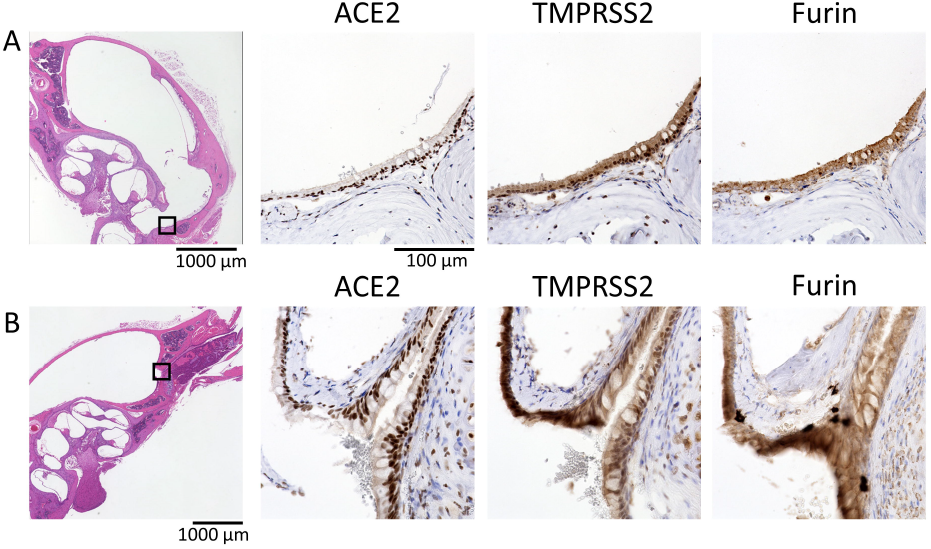
Histological analysis of the middle ear spaces and the Eustachian tube expressing Ace2, Tmprss2, and Furin. **A:** Hematoxylin-eosin staining of the middle ear mucosa (magnification, 40×). Immunohistochemical staining for Ace2 (cytoplasm), Tmprss2 (nucleus and cytoplasm), and Furin (cytoplasm) (400×). **B:** Parts of the Eustachian tube (40×) and Ace2, Tmprss2, and Furin expression in middle ear spaces (400×).

**Figure 2:**
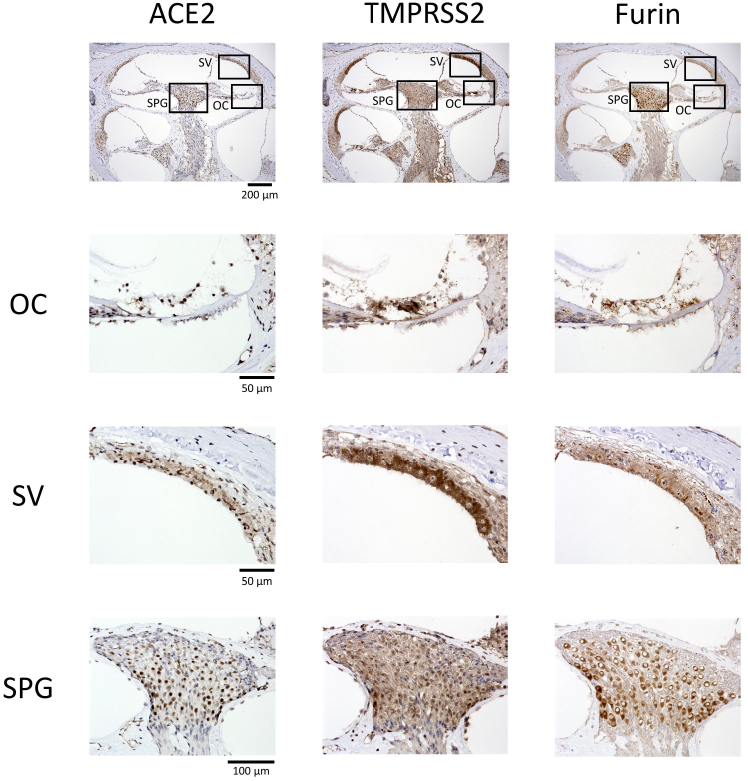
Cochlear Ace2, Tmprss2, and Furin expression. **Top panel:** Parts of the organ of Corti (OC), stria vascularis (SV), and spiral ganglion (SPG) (magnification, 40×). **OC**: Ace2 expressed in the nucleus of the hair cells and OC supporting cells; TMPRSS2, diffuse in OC cell nuclei and cytoplasm; Furin, diffuse in the OC cell cytoplasm. **SV**: Ace2, nucleus of marginal, intermediate, and basal cells in the stria vascularis; Tmprss2, nucleus and cytoplasm in the stria vascularis; Furin, cytoplasm in the stria vascularis. **SPG**: Ace2, nucleus of the spiral ganglion cells; Tmprss2, diffuse in the nucleus and cytoplasm in spiral ganglion cells; Furin, diffuse in the cytoplasm in the spiral ganglion cells.

### Histological analyses

Immunohistochemical staining was performed for Ace2 and Tmprss2. Four-micrometer-thick serial paraffin-embedded sections were deparaffinized in xylene and dehydrated in ethanol and then treated with 3% H2O2 to block endogenous peroxidase activity and incubated with Blocking One (Nacalai Tesque, 34 Kyoto, Japan) to block non-specific immunoglobulin binding prior to immunostaining. After antigen activation, the tissue sections were probed with primary anti-Ace2 (1:300 dilution; rabbit monoclonal, Abcam, ab108252; Cambridge, UK), anti-Tmprss2 (1:1000 dilution; rabbit monoclonal, Abcam, ab92323; Cambridge, UK), and anti-Furin (1:100 dilution; rabbit monoclonal, 1 Abcam, ab183495; Cambridge, UK) antibodies, followed by probing with appropriate peroxidase-conjugated secondary antibodies and a diaminobenzidine substrate. Images of all sections were captured using a digital microscope camera (Keyence BZ-X700) with 4×, 10×, and 40× objective lenses.

## Results

Ace2, Tmprss2, and Furin were detected in the mucosal epithelium of the Eustachian tube and middle ear spaces and in the cochlea, although their expression pattern varied among tissues. Furthermore, Ace2, Tmprss2, and Furin were co-expressed in the mucosal epithelium of the Eustachian tube and the middle ear spaces, the stria vascularis, and spiral ganglion cells.

Tmprss2 and Furin were expressed in the mucosal epithelial cells in the middle ear spaces; however, Ace2 was only slightly expressed in the cytoplasm of the epithelial cells (Fig. 1A). Furthermore, Ace2 and Tmprss2 were expressed in the nuclei of epithelial cells. Ace2, Tmprss2, and Furin expression profiles in the Eustachian tube were almost identical to those in the middle ear spaces (Fig. 1B).

In the cochlea, Ace2 was expressed in the nuclei of the hair cells and supporting cells in the organ of Corti, marginal, intermediate, and basal cells in the stria vascularis, fibrocytes of the spiral ligament, and spiral ganglion cells. Furthermore, Ace2 was slightly expressed in the cytoplasm of strial cells and spiral ganglion cells. Tmprss2 was diffusely strongly expressed in the nuclei and cytoplasm in the organ of Corti, stria vascularis, spiral ligament, and spiral ganglion cells, being greater in the nucleus than in the cytoplasm. Furin was diffusely expressed in the cytoplasm, but not in the nucleus, in the organ of Corti, stria vascularis, spiral ligament, and spiral ganglion cells (Fig. 2).

## Discussion

This immunohistochemical study shows that Ace2, Tmprss2, and Furin are co-expressed in the mucosal epithelium of the Eustachian tube and middle ear spaces and the organ of Corti, lateral wall, and spiral ganglion cells in the cochlea. In both middle ear tissues and the cochlea, Ace2 is primarily expressed in the nucleus, while Tmprss2 is expressed in the nucleus and cytoplasm, and Furin was expressed primarily in the cytoplasm. These results suggest that middle ear spaces are highly susceptible to SARS-CoV-2 infections spreading from the upper respiratory tract through the Eustachian tube, indicating the possibility of severe sensorineural hearing loss upon intracellular entry of SARS-CoV-2 in the cochlea.

Herein, we used ICR mice to determine the distribution of Ace2, Tmprss2, and Furin in the ear tissues. We could not obtain human ear tissue samples for immunostaining; hence, the expression patterns of these proteins in human ear tissues might differ from those of the present mouse ear tissues. However, we previously compared the Ace2, Tmprss2, and Furin expression patterns between human and mouse nasal tissues and found that their expression patterns were identical.^18^

Mastoidectomy is required during major ear surgeries including cochlear implantation and middle ear surgery for extensive cholesteatoma. Mastoidectomy with a high-speed drill generates massive aerosols.^19^ Previously, cadaveric simulation of otological procedures during mastoidectomy revealed gross contamination 3 to 6 feet away in all cardinal directions and more significantly on the left side, corresponding to the direction of drill rotation.^20^ SARS-CoV-2 would substantially infect the middle ear spaces because of the co-expression of Aces, Tmprss2, and Furin. Therefore, high levels of personal protective equipment are strongly recommended during mastoidectomy for COVID-19 patients or all mastoidectomies during the COVID-19 pandemic.

The cochlea is isolated and thus is generally resistant to the infection. A recent systemic review on the audio-vestibular symptoms of coronavirus infections found no records of audio-vestibular symptoms reported during previous coronaviral diseases including SARS and Middle East respiratory syndrome.^21^ However, Ace2, Tmprss2, and Furin were co-expressed in the organ of Corti, lateral wall, and spiral ganglion cells in the cochlea. Thus, similar to other viral infections including mumps, when SARS-CoV-2 enters the cochlea, it can cause severe labyrinthitis, resulting in marked hearing loss.

## Conclusions

Ace2, Tmprss2, and Furin are co-expressed in the mucosal epithelium of the Eustachian tube and middle ear spaces and the organ of Corti, lateral wall, and spiral ganglion cells in the cochlea in mice. ACE2 is primarily expressed in the nucleus, Tmprss2 is expressed in the nucleus and cytoplasm, and Furin is primarily expressed in the cytoplasm. These results indicate that middle ear spaces are highly susceptible to the SARS-CoV-2 infection, thus warranting the extensive use of personal protective equipment during mastoidectomy for COVID-19 patients.

## Acknowledgments

We would like to thank Editage (www.editage.com) for English language editing.

## Notes

**Funding:** This work was supported by JSPS KAKENHI Grant-in-Aid for Scientific Research (A) [grant number 20H00546].

**Conflict of Interest Statement**: We declare no competing interests.

### Competing Interest Statement

The authors have declared no competing interest.

